# Resting State Brain Connectivity analysis from EEG and FNIRS signals

**DOI:** 10.1101/2023.04.11.536366

**Authors:** Rosmary Blanco, Cemal Koba, Alessandro Crimi

**Affiliations:** Sano Centre for Computational Medicine, Computer Vision Data Science Team, Krakow, Poland

**Keywords:** EEG, fNIRS, multimodal monitoring, Functional Connectivity, Source-space analysis

## Abstract

Contemporary neuroscience is highly focused on the synergistic use of machine learning and network analysis. Indeed, network neuroscience analysis intensively capitalizes on clustering metrics and statistical tools. In this context, the integrated analysis of functional near-infrared spectroscopy (fNIRS) and electroencephalography (EEG) provides complementary information about the electrical and hemodynamic activity of the brain. Evidence supports the mechanism of the neurovascular coupling mediates brain processing. However, it is not well understood how the specific patterns of neuronal activity are represented by these techniques. Here we have investigated the topological properties of functional networks of the resting-state brain between synchronous EEG and fNIRS connectomes, across frequency bands, using source space analysis, and through graph theoretical approaches. We observed that at global-level analysis small-world topology network features for both modalities. The edge-wise analysis pointed out increased inter-hemispheric connectivity for oxy-hemoglobin compared to EEG, with no differences across the frequency bands. Our results show that graph features extracted from fNIRS can reflect both shortand longrange organization of neural activity, and that is able to characterize the large-scale network in the resting state. Further development of integrated analyses of the two modalities is required to fully benefit from the added value of each modality. However, the present study highlights that multimodal source space analysis approaches can be adopted to study brain functioning in healthy resting states, thus serving as a foundation for future work during tasks and in pathology, with the possibility of obtaining novel comprehensive biomarkers for neurological diseases.

## 1 Introduction

Large-scale functional brain connectivity can be modeled as a network or graph. For example, the synergistic use of cluster-based thresholding within statistical parametric maps could be useful to identify crucial connections in a graph. In the last few years, multimodal monitoring is getting increased interest. Integration of functional near-infrared spectroscopy (fNIRS) and electroencephalography (EEG) can reveal more comprehensive information associated with brain activity, taking advantage of their non-invasiveness, low cost, and portability. Between the two modalities, fNIRS relies on differential measurements of the backscattered light, which is sensitive to oxy(HbO) and deoxy-hemoglobin (HbR). EEG, on the other hand, captures the electrical brain activity scalp derived from synchronous post-synaptic potentials. The former has high spatial resolution but is highly sensitive to scalp-related (extracerebral) hemoglobin oscillations. The latter allows the tracking of the cerebral dynamics with the temporal detail of the neuronal processes (1 ms) but suffers from volume conduction. Their integration can compensate for their shortcomings and take advantage of their strengths [7, 6]. Most concurrent EEG-fNIRS studies focused on the temporal correlation of the time series data between these modalities. However, the two techniques do not have perfect spatiotemporal correspondence, since brain electrical activity and its hemodynamic counterpart are mediated through the neurovascular coupling mechanism. Therefore, comparing the correspondence between the two modalities from a more analytical and standardized perspective can be more informative. In this context, network neuroscience could be an approach to model the actual brain function and study the potential of multimodal approaches in inferring brain functions. To the best of our knowledge, little is known about the fNIRS-based functional connectivity related to EEG functional connectivity of large-scale networks from a graph-theoretical point of view. Therefore, this study aims to explore the topology of brain networks captured by the two modalities across neural oscillatory frequency bands in the resting state (RS) through graph theoretical approaches.

## 2 Materials and Methods

Synchronous resting state EEG and fNIRS recordings of healthy adults (n = 29; 28.5 ± 3.7 years) were obtained from an open dataset for hybrid brain-computer interfaces (BCIs) [23]. The recordings feature a 1 minute long pre-experiment resting-state data, an experimental paradigm including a motor imagery and a mental arithmetic test. As the focus of this work is RS only, we only processed and examined the RS part of the dataset. The 1-minute RS duration is justified for fNIRS as it was shown that the reliability of the network metrics stabilizes in 1 minute of scan [17, 13]. EEG data were recorded with 32 electrodes placed according to the international 10-5 system (AFp1, AFp2, AFF1h, AFF2h, AFF5h, AFF6h, F3, F4, F7, F8, FCC3h, FCC4h, FCC5h, FCC6h, T7, T8, Cz, CCP3h, CCP4h, CCP5h, CCP6h, Pz, P3, P4, P7, P8, PPO1h, PPO2h, POO1, POO2 and Fz for ground electrode). Those are referenced to the linked mastoid at 1000 Hz sampling rate (downsampled to 200 Hz). fNIRS data were collected by 36 channels (14 sources and 16 detectors with an inter-optode distance of 30 mm), following the standardized 10-20 EEG system, at 12.5 Hz sampling rate (downsampled to 10 Hz) as depicted in Figure 1. Two wavelengths at 760 nm and 850 nm were used to measure the changes in oxygenation levels. All the steps of the pipeline from EEG and fNIRS recording to the brain network are summarized in Figure 2.

**Fig. 1.**
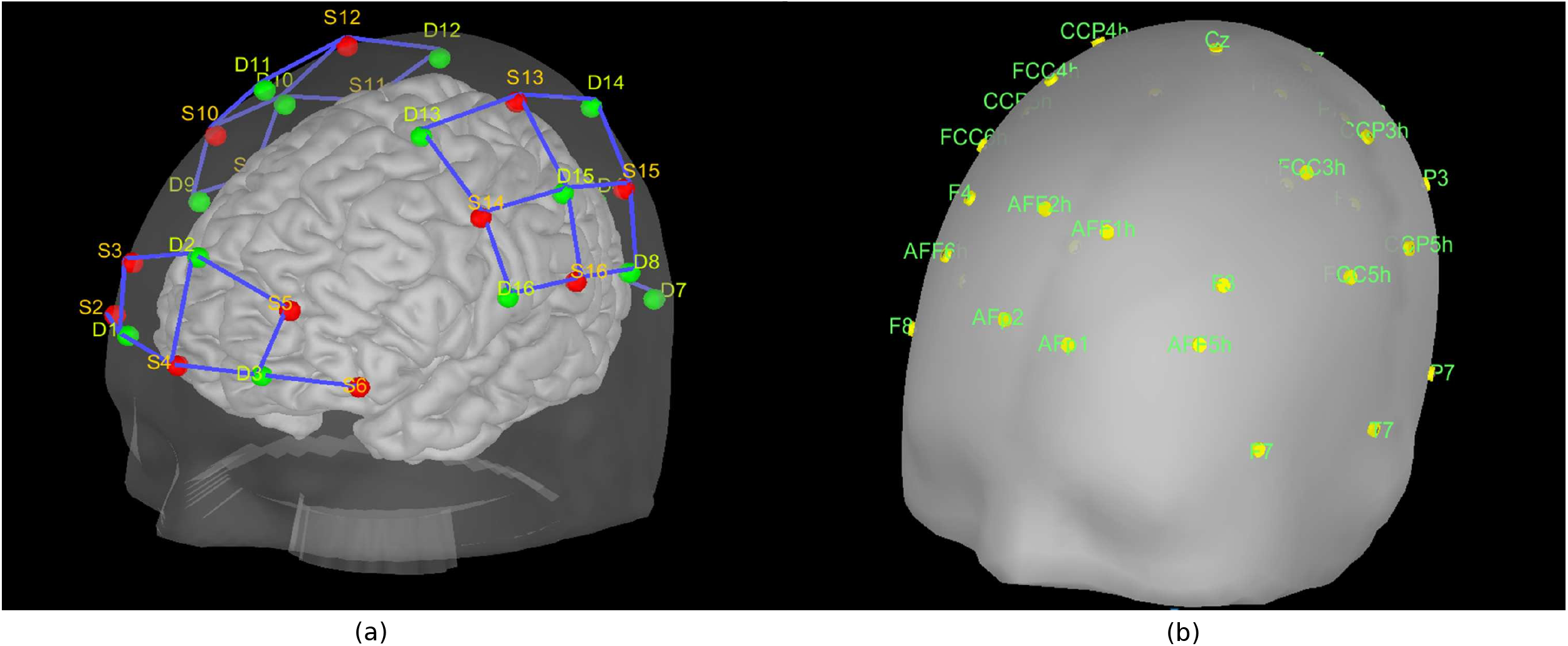
(a): NIRS optodes location. The red dots are the sources and the green dots are the detectors. (b): EEG electrodes location.

**Fig. 2.**
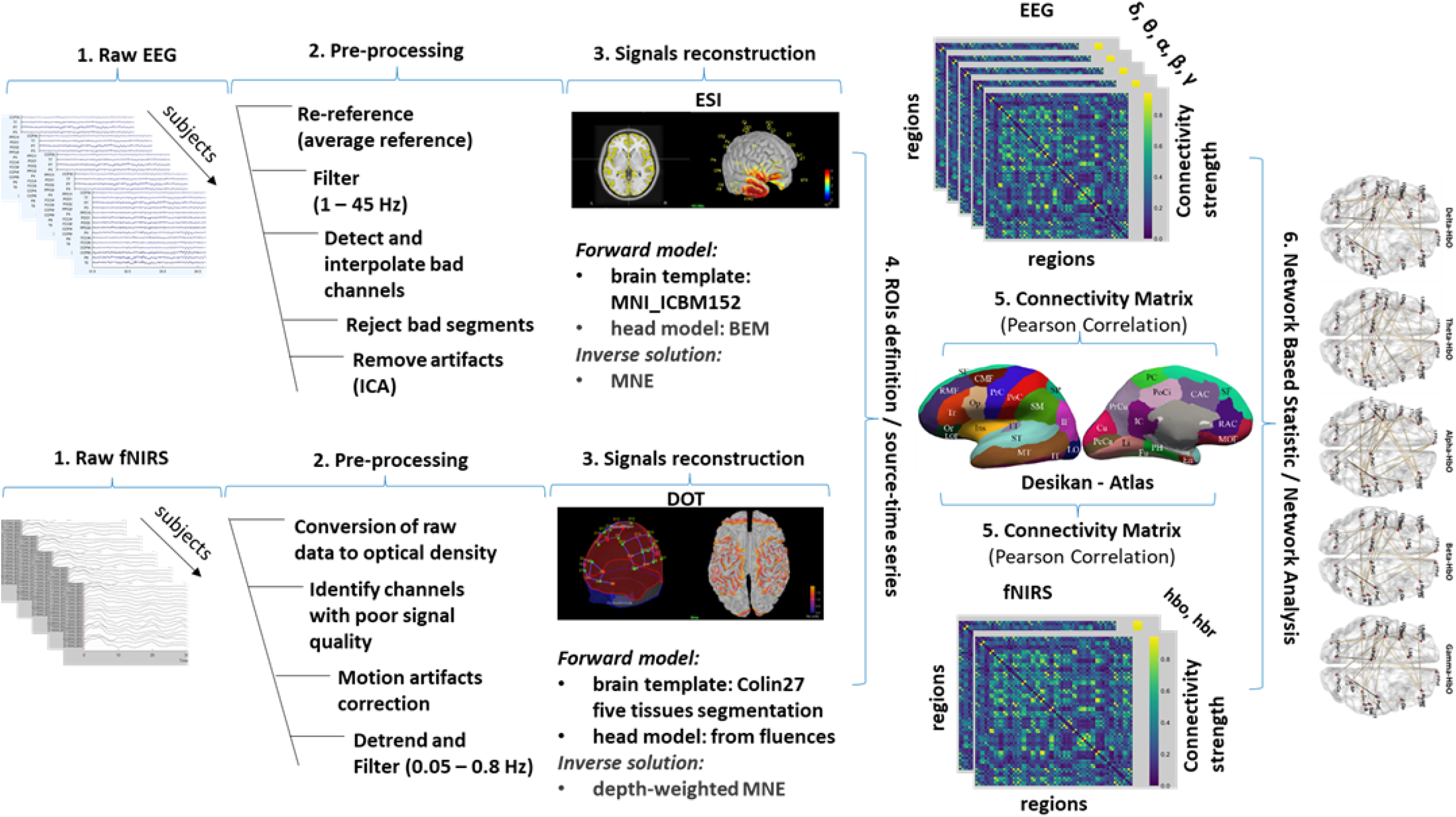
An overview of the entire pipeline from pre-processing to brain networks for both EEG and fNIRS data. 1. The EEG and fNIRS data recordings for all the subjects. 2. The pre-processing steps for cleaning data. 3. The methods for the reconstruction of the signals in the source space. The Electrical Source Imagine (ESI) technique aims to build a realistic head model (forward model) by segmenting the MRI to estimate the cortical sources (inverse problem). The result of the forward modeling (the lead field matrix) is the estimation of the contribution of each cortical source (neuronal activity) to the scalp sensors (differences in electrical potentials recorded by the electrodes). From the lead field matrix, the source model is built to estimate the amplitude of the current dipoles (solving the inverse problem). The Diffuse Optical Tomography (DOT) technique aims to model light propagation within the head by segmenting it into five tissue types and estimating the optical properties of each tissue by fluences of light for each optode (source and detector). The forward model is calculated from fluences by projecting the sensitivity values within the gray matter volume onto the cortical surface. The result of the modeling is the sensitivity matrix that maps the absorption changes along the cortical region (source space) to the scalp sensors (measured as optical density changes). The cortical changes of hemodynamic activity within the brain are estimated by inverting the forward model (solving the inverse problem). 4. The EEG and fNIRS source-time series were mapped in the same 3D space using an atlas-based approach (Desikan-Killiany) with the definition of Region of Interest (ROIs) at which both signals were estimated. 5. The statistical coupling between the reconstructed time series was estimated by the calculation of the functional connectivity using the Pearson correlation metric for both modalities. 6. The topology of brain networks captured by the two techniques was compared through graph theoretical approaches. As shown later in Figure 4.

### 2.1 EEG data pre-processing

The electrical source imaging (ESI) technique was used to estimate the cortical EEG activity in the source space [9]. A standardized pre-processing pipeline was applied to remove the artifacts prior to applying ESI using the EEGLab toolbox [4]. The measured EEG data was first re-referenced using a common average reference and filtered (second-order zero-phase Butterworth type IIR filter) with a passband of 1 – 45 Hz. Bad channels were identified, rejected, and interpolated taking an average of the signal from surrounding electrodes. The signals were then visually inspected to detect and reject segments of data still containing large artifacts. A decomposition analysis by using the fast fixed-point ICA (FastICA) algorithm, was performed to identify and remove artifacts of biological origin from the recordings [15].

### 2.2 fNIRS data pre-processing

We used diffuse optical tomography (DOT) to reconstruct the signals in the source space. More specifically, a combination of the Brainstorm toolbox (i.e., NIRSTORM) and a custom MATLAB script was used. Before the reconstruction, the typical pre-processing pipeline of the fNIRS data was applied. The raw data were converted to optical density (OD-absorption) signals for both wavelengths. The bad channels were removed based on the following criteria: signals had some negative values, flat signals (variance close to 0), and signals had too many flat segments. Then a semi-automatic movement correction was applied: the signals were visually checked for motion artifacts in the form of spikes or baseline shifts and a spline interpolation method was used for the correction. Finally, the OD signals were detrended and band-pass filtered (third-order zerophase Butterworth IIR filter) with a 0.05–0.8 Hz bandpass.

### 2.3 Signals reconstruction

Source EEG and fNIRS data reconstruction were performed using Brainstorm software [24] and custom Matlab scripts [2]. For EEG a multiple-layer head model (Boundary Element Method-BEM) and an MRI template (MNI-ICBM152) were used to build a realistic head model (forward model) by the OpenMEEG tool, which takes into account the different geometry and conductivity characteristics of the head tissues. The dipoles corresponding to potential brain sources were mapped to the cortical surface parcellated with a high-resolution mesh (15000 vertices). Finally, a leadfield matrix, expressing the scalp potentials corresponding to each single-dipole source configuration, was generated based on the volume conduction model [12].

For the fNIRS, a five tissues segmentation of the Colin27 brain template was used for the sensitivity matrix (forward model) computation. The fluences of light for each optode were estimated using the Monte Carlo simulations with a number of photons equal to 10^8^ and projecting the sensitivity values within each vertex of the cortex mesh. The sensitivity values in each voxel of each channel were computed by multiplication of fluence rate distributions using the adjoint method according to the Rytov approximation [22]. Then the NIRS forward model was computed from fluences by projecting the sensitivity values within each vertex of the cortex mesh using the Voronoi-based method, a volume-to-surface interpolation method which preserves sulci-gyral morphology[10].

The inverse problem (i.e., estimation of the source activity given a lead field matrix) was solved by the minimum norm estimate (MNE) method. The MN estimator is of the form:

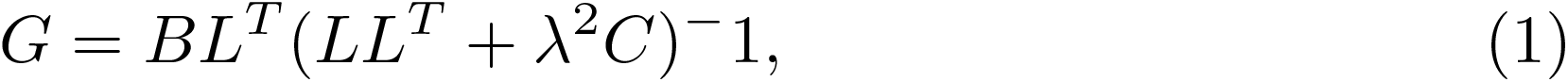

where G is the reconstructed signal along the cortical surface, L is the non-zero element matrix inversely proportional to the norm of the lead field vectors, B is the diagonal of L, C is the noise covariance matrix, and *λ* is the regularization parameter that sets the balance between reproduction of measured data and suppression of noise. For EEG the standardized low-resolution distributed imaging technique (sLORETA) was applied. While for fNIRS the depth-weighted minimum norm estimate (depth-weighted MNE) method was utilized. This was necessary since standard MNE tends to bias the inverse solution toward the more superficial generators, and the light sensitivity values decrease exponentially with depth.

In order to be comparable, the EEG and fNIRS source-time series were mapped in the same 3D space using an atlas-based approach (Desikan-Killiany) [8]. Since the optodes did not cover the entire scalp, it was necessary to modify the Desikan-Killiany atlas by choosing those ROIs covered by the fNIRS signal, leaving 42 ROIs out of 68 for both modalities. The EEG ROI time series were then decomposed into the typical oscillatory activity by band-pass filtering: *δ*(1 − 4*Hz*), *θ*(4 − 7*Hz*), *α*(8 − 15*Hz*), *β*(15 − 25*Hz*), and *γ*(25 − 45*Hz*), while fNIRS source reconstructions were converted into oxygenated hemoglobin (HbO) and deoxygenated hemoglobin (HbR) by applying the modified Beer-Lambert transformation [5].

### 2.4 Connectivity differences in brain networks

The functional connectivity matrices for the 42 ROIS within EEG (for each frequency band) and fNIRS (for HbO and HbR) were computed using Pearson’s correlation coefficient, generating 7 42 × 42 connectivity matrices for each subject (5 for each EEG frequency band and 2 for hemodynamic activity of fNRIS). Once the networks have been constructed, their topological features were calculated via *graph/network analysis* [19, 26]. More specifically, small-world index (SWI), global efficiency (GE), clustering coefficient (CC), and characteristic path length (PL) were chosen as the network features of interest.

The small-world topology of a network describes how efficient and costeffective the network is. A network is said to be a small-world network if SWI *>* 1. Brain networks with large small-world values are densely locally clustered, and at the same time employ the optimal number of distant connections, in this way the information processing is more efficient with lower information cost. Here the SWI was calculated as 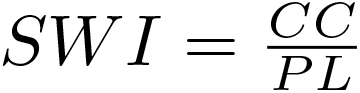.

As a measure of functional integration, the GE is defined as the efficiency of information exchange in a parallel system, in which all nodes are able of exchanging information via the shortest paths simultaneously.

The average shortest path length between all pairs of nodes is known as the PL of the network. As a measure of functional segregation, the CC is the fraction of triangles around an individual node reflecting, on average, the prevalence of clustered connectivity around individual nodes. The graph measures were normalized by generating 100 randomized networks (null models of a given network) and preserving the same number of nodes, edges, and degree distribution. Then, for each measure the ratio was calculated as *real metric* over the *matched random metrics*. For the comparison of network topology features, a paired student t-test was used between all the EEG frequency bands and the matched fNIRS metric of [(*δ, θ, α, β, γ* - HbO), (*δ, θ, α, β, γ* - HbR)], with the significance level set at p *<* 0.05. Multiple comparison correction was carried out using False Discovery Rate (FDR).

For the *edge-wise analysis*, that compare the connection strength between two regions, the network-based statistic (NBS) method was applied using the python brain connectivity toolbox [26]. For each edge of the 42 × 42 connectivity matrix, two-sample paired t-test was performed independently between each modality (5 frequency bands and 2 hemodynamic responses) and cluster-forming thresholds were applied to form a set of suprathreshold edges. The threshold was chosen based on Hedge’s g-statistic effect size (ES) computed between each node of the two matrices, pairing each EEG frequency band FC with each fNIRS FC. The t-statistic corresponding to the Hedge’s g score equal to 0.5 (medium ES) was chosen as the critical value (t-stat = 3.0). Finally, an FWER-corrected p-value was ascribed to each component through permutation testing (5,000 permutations). Edges that displayed FWER-corrected p-values below the significance threshold of 0.05 were considered positive results.

The topological characteristics of the graph, at the global and edge level, were calculated using the Matlab Brain Connectivity Toolbox and custom Matlab scripts [19].

## 3 Results

Topology analyses showed that, all the EEG frequency bands and fNIRS (HbO and HbR) have an SWI *>* 1, which implies prominent small-world properties. A real network would be considered small world if CC_*real*_/CC_*rand*_ *>* 1 and PL_*real*_/PL_*rand*_ *≈*1. It means that, compared to random networks, a true human brain network has a larger CC and an approximately identical PL between any pair of nodes in the network. This was demonstrated for all EEG frequency bands, showing significantly higher CC values than HbO, particularly for the lower frequency (*δ, θ, α*), associated with PL values around 1. That means better clustering ability and small-worldness than HbO. As for HbR, the clustering coefficients were higher with respect to all EEG frequency bands (Figure 3) Also, significantly higher E values in *δ, θ, α, β, γ* vs. HbO and HbR, were pointed out. (Figure 4). This means that, for electrical brain activity, the neural information is routed via more globally optimal and shortest paths compared to hemodynamic activity. Thus, they provide faster and more direct information transfer.

**Fig. 3.**
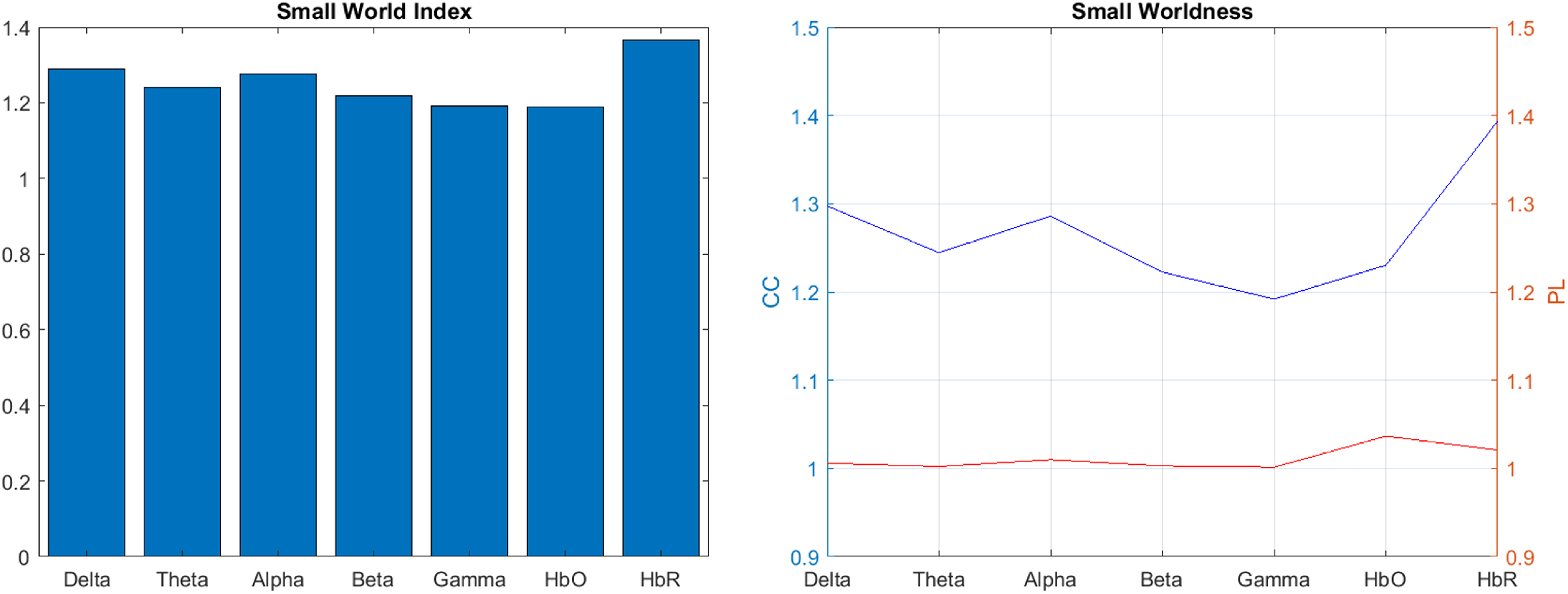
Left: barplot of Small-World Index (SWI), Right: plot of Global Characteristic Path Length (PL) and Global Clustering Coefficient (CC) for EEG (Delta, Theta, Alpha, Beta, Gamma) and fNIRS (HbO and HbR)

**Fig. 4.**
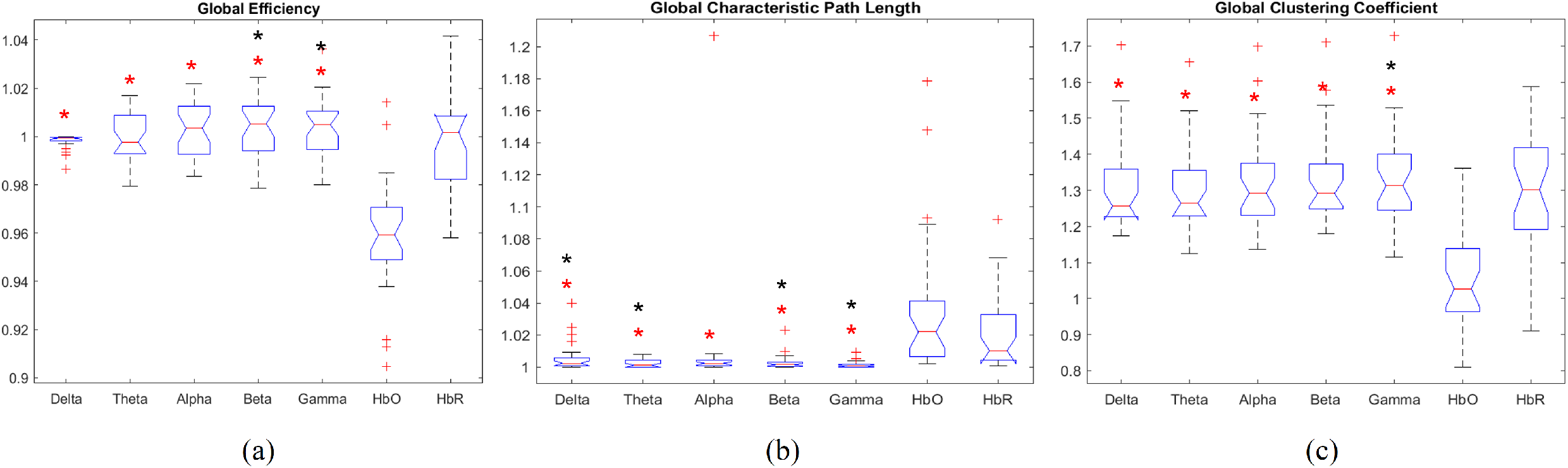
Boxplot of (a) Global Efficiency (E), (b) Global Characteristic Path Length (PL) and (c) Global Clustering Coefficient (CC) for EEG (Delta, Theta, Alpha, Beta, Gamma) and fNIRS (HbO and HbR). The red asterisk denotes the significant difference between EEG and fNIRS (HbO); The black asterisk denotes the significant difference between EEG and fNIRS (HbR).

**Fig. 5.**
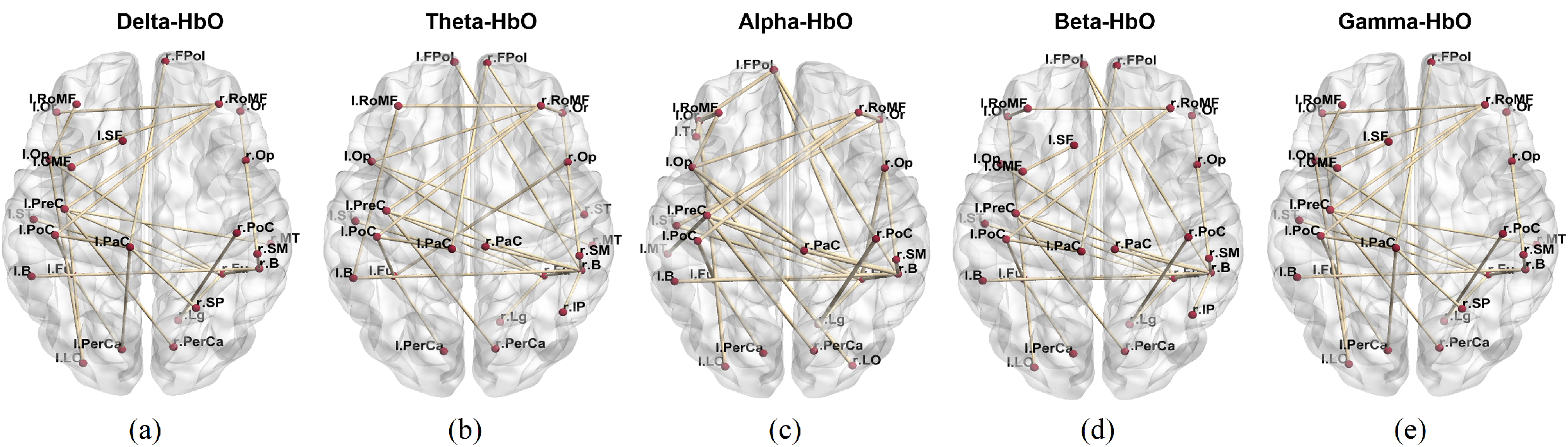
Subnetwork between each pair of fNIRS (HbO) and EEG frequency band: (a) Delta, (b) Theta, (c) Alpha, (d) Beta, (e) Gamma. Red dots denote the nodes and the yellow links the edges of the network.

At edge-wise analysis, the NBS identified one subnetwork of increased functional connectivity for HbO compared to EEG for all the frequency bands [*δ* (p=0.005), *θ* (p=0.005), *α* (p=0.004), *β* (p=0.004), *γ* (p=0.003), Figure 3, corrected for multiple comparisons] at a pre-defined threshold of 3.5. This subnetwork consisted of 51 edges for *δ*-HbO connecting 25 nodes, 57 for *θ*-HbO connecting 23 nodes, 63 for *α*-HbO connecting 24 nodes, 59 for *β*-HbO connecting 26 nodes, and 57 for *γ*-HbO connecting 25 nodes.

These subnetworks, similar across all frequency bands, are characterized by inter-hemispheric and intra-hemispheric connections. The former comprises connections between: 1- the right superior and medial frontal gyrus at the premotor area (FPol, RoMF) with both the left superior frontal sulcus, at the human frontal eye field (Op, Or, Tr) and left pre- and post-central gyrus at the primary sensory-motor cortex (PreC, PaC, PoC); 2- the left sensory-motor cortex (PreC, PaC, PoC) with both the right temporal-occipital regions (MT, B, Fu) and right posterior cingulate/precuneus (PerCa, LO) gyrus; 3- the medial frontal gyrus (RoMF) and the temporo-occipital cortex (B) with their homotopic regions. The latter comprises connections between: 1- the right frontal eye field (Op, Or, Tr) with right temporal-occipital regions (B); 2- right B with the right lingual gyrus (Lg); 3right postcentral gyrus (PoC) with right Lg. 4- Left superior and median frontal sulcus (Or, RoMF) with left posterior precuneus (LO); 5- left post-central gyrus (PoC), with left pericalcarine (PerCa) regions. The full list of the labels from the atlas is given in [8].

## 4 Discussion

To the best of our knowledge, we are the first to investigate the topological properties of functional networks between synchronous EEG and fNIRS connectomes, across frequency bands, using source space analysis. We have observed a small-world topology network for both modalities, suggesting that the smallworldness is a universal principle for the functional wiring of the human brain regardless of the distinct mechanisms of different imaging techniques. The brain supports both segregated and distributed information processing, a key for cognitive processing, which means that localized activity in specialized regions is spread by coherent oscillations in large-scale distributed systems [19]. Our results have shown a significantly lower CC and large PL for HbO than EEG, indicating a lower specialization ability and a lower ability for parallel information transmission in the HbO network. The network differences between HbO-derived hemodynamic activity and EEG mostly pointed out inter-hemispheric connections, and to a lesser extent intra-hemispheric connections, between prefrontal and temporo-occipital regions. In agreement with the previous study [20], which reported HbO differences during RS in both short-distance ipsilateral connectivity between the prefrontal and occipital regions, and long-distance contralateral between homologous cortical regions. It was suggested that the generation of homologous connectivity is through direct structural connection, while fronto-posterior connectivity may reflect the synchronization of transient neural activation among distant cortical regions [20]. The involvement of different oscillatory frequencies in a cortical network varies as a function of cognitive state [3], and those states often last only a few ms. Since it is accepted that the shortand long-range organization of neural activity by oscillations of different frequencies depends on how those oscillations work in concert and that more than one EEG frequency band has been associated with the same RS network, our data may reflect unknown state changes in RS [14]. Thus, given that the hemodynamic response is delayed by several seconds, fNIRS cannot distinguish between these rapidly changing neural responses, reflecting the sum of several oscillatory network configurations.

The EEG activity was associated with metabolic deactivation in numerous studies. Moosmann et al. (2003) [16] measuring simultaneous EEG-fNIRS RS, found that the alpha activity had a positive correlation with HbR in the occipital cortex. While Koch et al. 2008 [11] reported that a high individual alpha frequency (IAF) peak correlates with a low oxygenation response. They conjectured that the relationship between IAF, neuronal and vascular response depended on the size of the recruited neuronal population. Another possible explanation is that since the oxy-hemoglobin is closely related to local cerebral blood flows, an RS condition correlates with lower metabolic demand [25]. However, it is accepted that functional connectivity maps derived from hemoglobin concentration changes reflect both spontaneous neural activity and systemic physiological contribution. Roughly 94% of the signal measured by a regular fNIRS channel (source-detector distances of 3 cm) reflects these systemic hemodynamic changes, producing low-frequency fluctuations. Thus contributing to a higher proportion of the variance of the HbO signal and to highly correlated brain regions, even after motion artifact correction and pre-whitening were employed, overestimating RS FC [1]. Furthermore, it was observed that oxy-hemoglobin is more heavily contaminated by extracerebral physiology than deoxy-hemoglobin, which has less sensitivity to systemic physiology [21]. To conclude, the results of this study point to the characteristic differences between electrophysiological and hemodynamic networks. Their simultaneous use is suggested since they can give a clearer picture of brain dynamics. However, it is crucial to understand the characteristics of each modality and understand what the differences and similarities mean to interpret them correctly. In this context, combining the brain network topological metrics based on graph theory analysis with multiple machine learning (ML) algorithms could be used to extract significant discriminative features, which could help to reveal the changes in the underlying brain network topological properties. It will be also interesting to study if those graph features will be able to capture transient and localized neuronal activity during task conditions. Future studies will investigate how the technological differences between EEG and fNIRS, when associated with ML, could be useful to extract discriminating features in pathology such as Alzheimer’s disease.

## Acknowledgements

This research study was conducted retrospectively using human subject data made available in open access by [23]. Ethical approval was not required as confirmed by the license attached with the open access data.

This research is supported by the European Union’s Horizon 2020 research and innovation programme under grant agreement Sano no 857533, and by the International Research Agendas programme of the Foundation for Polish Science, co-financed by the European Union under the European Regional Development Fund. This research was tested on D-Wave quantum computing.

